# Night temperature determines nearly half of wheat yield variation globally

**DOI:** 10.64898/2025.12.19.695361

**Authors:** Urs Schulthess, Matthew Paul Reynolds, Owen Kenneth Atkin, Ernesto Giron, Senthold Asseng, Sieglinde Snapp

## Abstract

Daily minimum temperature (Tmin) is increasing faster than maximum temperature (Tmax). However, the impact of Tmin on crop productivity is barely studied. Effects of environmental covariates on yield were examined using 42 years of trials at 255 sites representing spring wheat regions globally. Grain-filling was the most sensitive growth stage: Average Tmin explained 40% of yield variation, and 52% when also considering radiation, while average Tmax explained just 20% of variation. Yield declined linearly between the observed range from 8 to 22ºC average Tmin. Generally, a 1ºC increased Tmin reduced yield by ∼0.5 t/ha, with high radiation partially offsetting negative effects. Average increase of 1.2ºC at test sites over 42 years reduced yield by more than 10%. Shorter grain-filling duration likely reduced yield, as well as increased nocturnal rate of dark respiration. A better understanding of drivers of variation in respiration and adaptation to warmer nights could generate a step change in wheat yield.

## Introduction

Climate change has increased the frequency and intensity of hot weather events^1^, and led to a steep increase in nighttime minimum (Tmin) relative to daytime maximum (Tmax) temperature gains^2^. Importantly, the effects of global warming differ by region^3^, with a review paper by Ullah et al.^4^ concluding that the effects of extreme temperatures on grain yield of wheat are variable; hence, one cannot expect identical changes in temperatures across all wheat production regions. One contributing factor is that irrigated wheat can cool its canopy up to 8ºC during the day in semi-arid regions^5^, partially mitigating the effects of high daytime temperatures. To quantify the effect of global warming on historic and future wheat yields, crop simulation models take weather, as well as soil and cultivar characteristics into account^6,7,8^. Assessing the plausibility of the simulated impact of climate change on wheat production remains challenging, as the simulation results depend on many inputs, such as climate scenarios, and the response of wheat to warming nighttime temperatures is still not fully understood. This is of major concern as crop productivity is negatively coupled to night temperature^9,10^, for yet unknown reasons. Even basic research has not been able to discern the mechanisms by which increasing nighttime temperatures reduce yield^11^, or interaction with growth stage. One of the challenges in understanding the effect of warming nights is that field research tends to show show different results than detailed studies conducted under controlled conditions^12^.

Wheat evolved in the Fertile Crescent, where it is exposed to cool temperatures during the winter months and high daytime temperatures, as well as dry and sunny conditions during the grain filling period, which coincides with the end of the rainy season in the spring. During the roughly three week period from heading, which coincides with the end of leaf growth, until the beginning of grain filling, organ growth is minimal and assimilates are stored in the form of water soluble carbohydrates and starch, to be subsequently remobilized for grain growth and maintenance respiration, especially under drought conditions^13–15^. This capacity to buffer adverse environmental conditions makes it challenging to quantify the effects of temperature and solar radiation on grain yield. As Evans et al.^16^ point out, measured responses to heat can change over time and across scales, i.e., from biochemical kinetics to biome effects. For example, thermal acclimation of dark respiration to sustained rises in nighttime temperatures can greatly alter the net biomass accumulation, leading to discrepancies between results obtained under artificial conditions and in the field^17–19^.

CIMMYT has sent advanced nursery lines for testing by partners across all major wheat production regions on Earth since the 1970’s. In return, partners share back key observations and measurements, such as yield. The lack of daily weather data has limited analyses until recently, whereby advances in reconstructing historic weather data to hourly and daily timesteps now allow for thorough analyses of the effects of weather on yield^20^. Based on a global data set spanning more than 40 years, the objective was to identify environmental variables with global impact on yield. Results enable identification of lines and genomes with adaptation to Tmin and other environmental conditions. This provides ‘model genotypes’ to guide research and is a diversity source for breeding.

## Results

Our data set consists of 850 site-years. It spans a period of 42 years and represents most of the important spring wheat growing areas on Earth (Fig.1). We tested the effects of temperature, solar radiation, and vapor pressure deficit on grain yield during three cardinal growth periods: 1) sowing to heading minus 10 days when the canopy is established; 2) heading +/-10 days when grain set is determined; and 3) grain filling. The analysis showed that weather conditions during the grain filling period have the strongest association with yield variation: average daily minimum temperature (Tmin) during this period ranked first, followed by average daily solar radiation (Supplementary Fig. 1). These two parameters were followed by the minimum temperature during the heading period. The prediction powers of average daily Tmin and Tmax, as well as solar radiation during grain filling for grain yield are shown in Supplementary Fig. 2.

The locations with the coolest nights or lowest minimum temperatures during the grain filling period are located in Egypt and Chile. These locations coincide with those that have the highest grain yields^21^. The Yaqui Valley in northwest Mexico also has high solar radiation, but slightly warmer Tmin. Sudan, Nigeria, and several countries in South Asia had the warmest nighttime Tmin. The lowest yields were recorded in Laos and Thailand, where warm nights coincide with low solar radiation during the grain filling period.

Combined Tmin and solar radiation during grain filling explains 52% of the variability of grain yield at sites across the globe, with a root mean square error of 1.38 t/ha (Fig. 2). Our data set which encompasses the majority of spring wheat production regions, shows that the yield response to average nighttime Tmin is linear within the observed range from 8 to 22ºC. An increase of 1ºC in average Tmin reduces yield by 0.5 t/ha. The results also show that high solar radiation can help plants compensate for warm nighttime Tmin: even under a warm average nighttime Tmin of e.g., 18ºC, yields increased from 2.6 t/ha to 5.9 t/ha when average daily solar radiation increased from 15 to 31 MJ/m^2^/day. Notably, the interaction between nighttime Tmin and solar radiation was not significant; hence, the effect of solar radiation is constant over the entire range of Tmin. The combination of low radiation and warm nighttime temperature causes the lowest yields. Vice versa, cool nights in high radiation environments, such as in Egypt or the Yaqui Valley in Mexico, result in the highest yields. The capacity of wheat to take advantage of high solar radiation levels can presumably be traced back to its origins in the Fertile Crescent, which helped it propagate under conditions of limited and erratic water supply combined with high temperatures and solar radiation during grain filling. This analysis shows that high solar radiation helps wheat tolerate warm nights.

There is a large regional variability in the increase in nighttime Tmin during grain filling over the period from 1980 to 2021 (Fig. 3). The largest increases were observed in the Mediterranean basin. They were generally above 1.5ºC. Central Europe, Sudan, Mesopotamia, and Western China experienced similar temperature increases. In the other parts of the world, the Tmin increases were generally below 1.5ºC. The lowest increases were observed for Chile, Argentina, and the Canadian Prairies, as well as north-east China. These are important spring wheat-producing regions. Interestingly, in India and Pakistan, the warming trend was higher at sites in the Indo-Gangetic Plain than at those further south.

Based on trends in Tmin changes, we calculated their impact on yield, using the estimated value of a yield loss of 0.5 t/ha for a 1ºC increase. The yield declines over the past 42 years, expressed in percent of the reported yields (Supplementary Fig. 3), tended to be larger in locations that are already low yielding, such as Sudan, where the average Tmin during the grain filling are higher than 20ºC (Fig. 4). In that country, the yield loss at three out of 4 locations exceeded more than 40%. Yield declines of more than 20% percent were estimated for Morocco, Central Europe, Mongolia, and five out of 19 locations in Pakistan. The majority of sites in Spain, Portugal, Iran, Syria, Türkiye, Afghanistan, and China had yield losses of 10 to 20%. In India, Iran, Mexico, and Türkiye, the yield losses hovered at around 10%. Slightly lower yield losses were observed for Brazil. In southern Africa, yield losses were below 10% in Zambia, Zimbabwe, and South Africa. Relatively small yield losses were also observed for Argentina, Chile, and Canada. It is noteworthy that in Chile, the country with the highest reported average yield, yield losses were close to zero (Supplementary Table 1).

## Discussion

The recent availability of globally gridded, reconstructed weather data (AgERA5) enabled us to disentangle environmental effects on yield over a 42-year period using measured plot-level data from 255 locations representative of most spring wheat production regions. The reasons for the finding that nighttime minimum temperatures (Tmin) during the grainfilling period control grain yield to a much larger extent than generally assumed are still not fully understood, especially since results from field studies differ from those obtained under controlled conditions^22^.

In a recent review, Amthor^11^ hypothesized that warming effects on truncating the duration of grain filling^23^, and thus grain yield, may override the effects of warming *per se* on specific rates of photosynthesis and respiration, i.e, yield declines are mostly due to a change in phenology. We had records of observed days to maturity for 262 site-years. As shown in Supplementary Fig. 4, grain filling duration indeed was shortened by warmer nighttime Tmin. Each increase in Tmin by one degree Celsius accounted for a shortening by 1.6 days. However, the prediction accuracy was not very high, with an R^2^ of 0.27 and RMSE of 6.4 days. In order to quantify the relative importance of the two factors, we calculated the Variable Importance Index, that reflects the relative contribution of a factor alone, not in combination with other factors. Using the dependent resampled inputs options, the report revealed that the main effect of nighttime Tmin on yield was 0.56, while it was 0.44 for duration of grain filling. Hence the shortened duration of grain filling is important. However, the direct effect of nighttime Tmin on other processes carried even more weight.

Due to the scale, it was not possible to measure photosynthesis and respiration; however, observed grain yield is a function of these two metabolic processes along with partition of assimilates. Daytime temperature has a strong effect on the rate of photosynthesis, i.e., the production of ATP and NADPH (energy and reducing power) needed to fix atmospheric CO_2_ and synthesize sugars like glucose, while nighttime temperatures primarily stimulate dark respiration rates^24,25^, the process that produces ATP for cellular maintenance and plant growth. It is noteworthy that higher dark respiration rates can create a larger sink for photosynthates, leading to higher photosynthesis rates under sink-limited conditions^24,25^. A key process that is likely accelerated by night warming is nocturnal protein turnover^22^ – which increases demand for respiratory ATP and the rate of respiratory CO_2_ release in darkness^22^. Another process is the effect of night warming on membrane fluidity, with negative consequences for the efficiency of respiratory ATP synthesis and the scale of nocturnal respiratory CO_2_ release^26,27^. The loss of carbon by respiration matters, as even under optimal conditions whole-plant respiration releases 30–60% of the carbon fixed by photosynthesis each day^28,29,30,31^ with approximately half of the CO_2_ released by whole-plant respiration coming from leaves during the night^32,33^. High rates of leaf respiration – a result of either increased demand for respiratory ATP to repair degraded proteins and/or decreased efficiency of respiratory ATP synthesis – during warm nights thus have the potential to play an important role in controlling daily carbon gain. This has consequences for both plant growth and grain yield.

Our study, at the biome level, helps to better understand how temperature and solar radiation affect these two metabolic processes: photosynthesis and nocturnal respiration. The fact that nighttime Tmin are more closely related to yield than daytime maximum temperatures emphasizes the important role of nocturnal dark respiration, a process which has received much less attention than photosynthesis. While both photosynthesis and respiration matter, growth rates of wild and crop plants are often much more strongly correlated with dark respiration than with photosynthesis^34^. The importance of respiration for growth and yield is supported by the finding that low-respiring ryegrass lines exhibited the highest yield^35^. Our study, which found that increasing night Tmin reduces yields, complements these findings and emphasizes the importance of nocturnal rates of dark respiration.

One factor that likely influences the extent to which warm-night mediated increases in respiratory CO_2_ release influence yield is whether respiration acclimates to sustained increases in night temperature. Acclimation to high growth temperatures often results in reduced rates of leaf dark respiration at any given temperature^36^, potentially reducing the stimulatory effect of warm nights on respiration rates. For wheat, this can result in homeostatic rates of whole-plant respiratory CO_2_ release in plants experiencing both warm days and nights, compared to those grown under cooler days/nights, when each (i.e. cool and warm grown plants) are measured at their respective growth temperatures^28^. However, in a recent study assessing the impact of warm nights *per se* on leaf respiration and growth in wheat, no acclimation of leaf respiratory CO_2_ release was observed^37^. As a result, warm nights stimulated respiratory CO_2_ loss and reduced plant growth. If similar patterns of minimal acclimation responses to warm nights occur at the field sites used in our study (particularly during grain filling), it would explain why warm nights were associated with reduced yields and partially compensated for by high solar radiation. In the future, improvements in yield under warm nights may be possible through screening for genotypic variability in thermal acclimation potential, building on the recent observations that leaf dark respiration is partially under genetic control^38^ and that there is genotypic variation in the extent of acclimation of wheat respiration to warmer nights^39^. A better understanding of dark respiration in combination with targeted breeding efforts may help limit yield losses caused by global warming, which creates a sharp increase in nighttime temperatures.

Our estimated yield losses tend to be larger than the forecasts generated by ensembles of crop simulation models in conjunction with climate change scenarios^7,8^. These two papers cover different periods: while the paper by Asseng et al.^7^ focused on hot spots of yield decline for irrigated spring wheat for the period from 2030 to 2041, Jägermeyr et al.^8^ covered most current and potential wheat production regions on Earth between 1980 and 2100. Unlike these studies, our study did not account for the contested effects of CO_2_ fertilization^40^. A recent meta-analysis^40^ concluded that rising temperatures can negate the effects of CO_2_ fertilization. Nevertheless, one would expect similar regional trends, even when our study focuses on the past. Asseng et al.^7^ forecast yield declines in all regions, which matches our analyses: Our studies agree on small yield declines for Egypt, and strong yield declines for Sudan, Saudi Arabia, and Western China. However, for India and Pakistan, our model estimates the largest relative yield losses for the regions with the highest production levels: Punjab in Pakistan and India, and Haryana in India, and smaller losses for the regions further south and east, which is the opposite pattern predicted by the multimodel ensemble. We also estimate relatively larger yield declines for Mesopotamia. The global study by Jägermeyr et al. agrees with our estimate of largest declines in Sudan and Saudi Arabia, and intermediate declines in Mexico. We estimate only small declines for Chile and Canada, whereas Jägermeyr estimates yield increases for these countries by 2100. The largest discrepancies between our nighttime temperature based model and Jägermeyr et al. are observed for the Mediterranean, the Middle East and Western China, for which we estimate a yield decline since 1980 by 10 to 20%, whereas Jägermeyr estimates a slight increase in yield by 2100 in the Mediterranean and a strong yield increase for the Middle East, Central Asia and all of China. While our model is empirical, based on daily minimum temperature and solar radiation along with measured yield data, crop simulation models are mainly driven by radiation use efficiency^41^ and rely on a number of assumptions.

Our observations show much bleaker trends for wheat production in these regions. Unlike the crop simulation models, we do not take rainfall into account. Given the fact that rainfall amounts have shown a declining trend and have become more erratic over the past four decades in Mesopotamia and Türkiye^42,43^, it is unrealistic to foresee an increase in wheat yields in this region, where wheat is the most important staple crop. This has serious negative implications for food security.

Our study highlights that warming trends vary greatly across wheat production regions. Notably, the Mediterranean basin and Mesopotamia are much more affected by global warming than those located south of the equator. Our results, which are representative for most of the important spring wheat production regions on Earth, also indicate that the effects of warming on grain yield of wheat are more severe than those forecasted by ensembles of crop simulation models or from meta-analyses^22,44^.

## Conclusions

Previous studies have mostly focused on the effects of maximum temperatures on crop yield, and some have shown a negative association between yield and Tmin in staples such as rice and maize, though growth stage was not considered. Our study, based on field-grown wheat representing most of the spring wheat production regions globally, demonstrates that increases in nighttime minimum, not daytime maximum temperatures, have limited the grain yield of wheat. This is partly due to a shortening of the grain filling period, but to a larger extent caused by the impact of warmer nights on yield – an observation that points to increased rates of respiratory carbon dioxide loss as a major factor driving reduced wheat yields across the globe. Hence, to increase yields, or rather, limit yield losses due to warming, there needs to be an increased focus on understanding drivers of variation in respiration of wheat. Our study also challenges the assumptions of existing crop simulation models, which suggest that variations in yield are mainly driven by photosynthesis. Hence, such models cannot properly quantify the impact of global warming, and especially the effect of the relatively sharper increase of nighttime, as compared to daytime maximum temperatures, on crop yield.

Our nighttime-based analysis shows an alarming trend, with the potential yield generally declining by 10% or more over the past four decades. More efforts and resources are needed to intensify breeding efforts for lines that better tolerate high nighttime temperatures by minimizing respiration under warm night conditions. These efforts need to be complemented by better agronomic practices, such as conservation agriculture, that improve soil health and can buffer soil moisture from weather anomalies where crop residue retention is practiced and thus foster yields^45,46^. Most of the ‘low-hanging fruits’ in crop improvement, such as improved harvest index, have been exploited. New breeding tools, such as genomic selection and gene editing, offer new paths to accelerate breeding gains, but we first need to better understand the mechanisms of how nighttime temperature and nighttime physiology in general control key physiological processes, including dark respiration.

Our results are of considerable importance to food security and commerce as well as updating crop simulation and breeding models. They fill a gap in fundamental research by showing that growth is tightly coupled to the wide range of night temperatures, under which wheat is currently grown. Hence, a significant improvement of adaptation to warmer nights has the potential to generate a step change in yield over the majority of current wheat-growing environments, which already experience supra-optimal night temperatures. Similarly, contributors to increased respiratory costs for cellular maintenance - which include protein turnover and maintenance of ion gradients across membranes - along with the energy costs of starch synthesis in growing spikes, will need to be studied for genetic variation and potential to improve efficiency. For example, starch synthase is well known to be heat-labile. Genetic variation in diurnal cycles and temperature thresholds may help to identify the genetic bases of components of maintenance respiration. Differences in varietal response to warming nighttime temperatures identified by this study can be used in crossing strategies almost immediately.

## Material and Methods

We used a data set derived from the Elite Spring Wheat Yield Trials (ESWYT) for this analysis. Under the umbrella of the International Wheat Improvement Network (IWIN), CIMMYT sends out different nurseries consisting of 30 to 50 advanced lines for testing by its partners across all the major wheat production regions on Earth (Fig. 1). In return for getting access to advanced genetic material, the partners record many parameters, such as sowing date, days to heading and maturity, plant height, disease incidence levels and yield. However, some of the returned data sets have missing values. Days to heading was available for 92% of the records, while days to maturity was reported for 32%. Missing phenological data were gap-filled. The data set was quality controlled, resulting in 850 site-year combinations. For each combination and line, we calculated the best linear unbiased estimation (BLUE) yield. Each nursery year consists mostly of new lines, and only a few are retained for more than one year. All in all, 1402 lines were tested. We extracted the daily weather parameters from AgERA5^20^ and calculated weather related parameters for the following three periods: 1) sowing to heading minus 10 days, 2) heading minus 10 days to heading plus 10 days, and 3) heading plus 10 days to maturity. The last period will be referred to as grain filling. The first period covers the establishment of the plants and tiller growth, the second one lasts from booting to the beginning of grain filling, the period around anthesis, during which organ growth is minimal, and the last one coincides with grain filling. For each period, we calculated average daily minimum and maximum temperature, solar radiation, vapor pressure deficit (VPD), number of days on which daily maximum temperature (Tmax) is exceeding 30ºC and daily minimum temperature (Tmin) is exceeding 25ºC, and vapor pressure deficit VPD exceeding 4 kPa. For ease of understanding, we will refer to daily minimum temperature as nighttime temperature, although they are not precisely the same.

**Figure 1:**
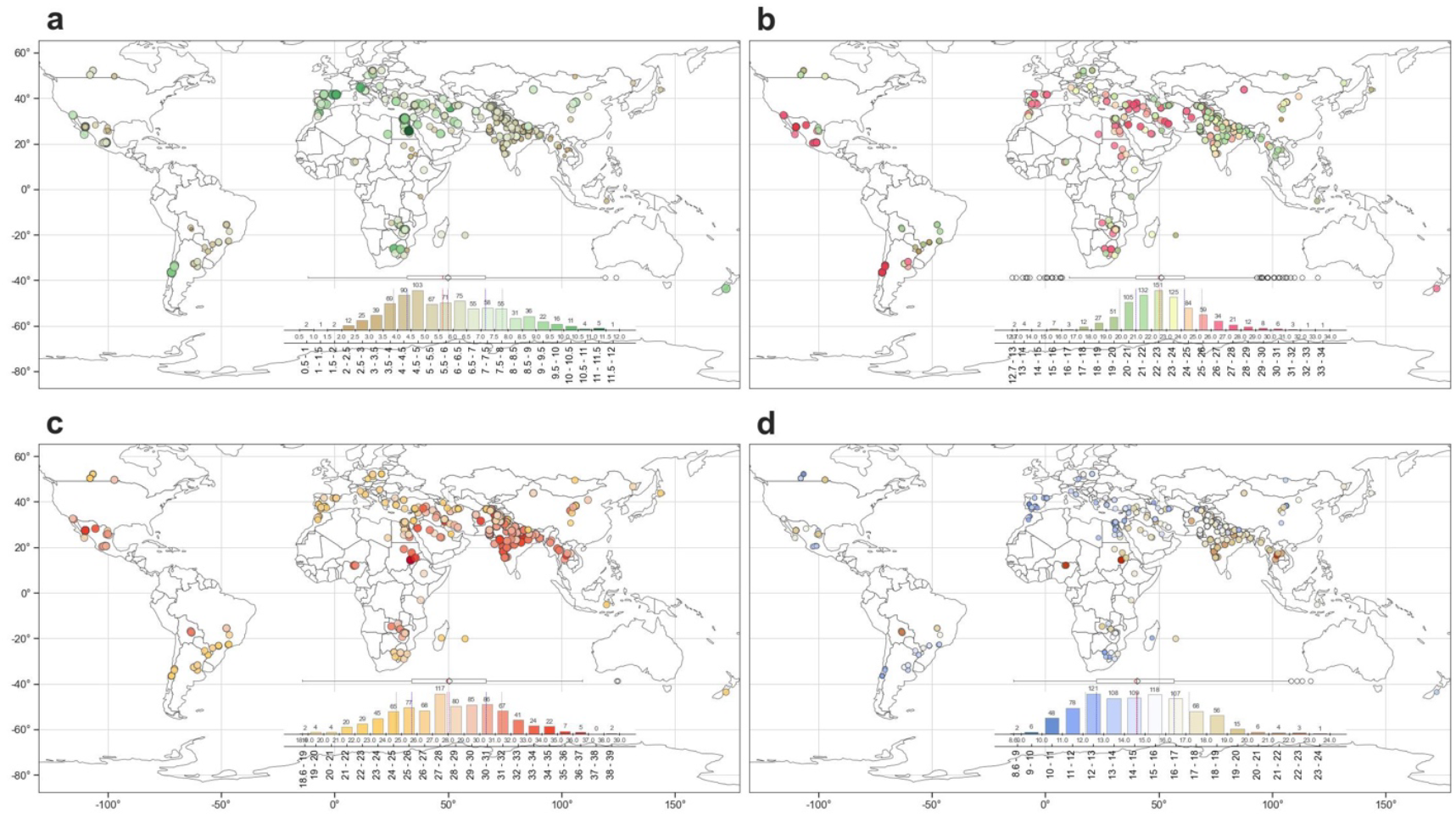
**a)** Grain yield, and **b)** average daily solar radiation, **c)** maximum and **d)** minimum temperatures during grain filling at 255 test sites for the Elite Spring Wheat Yield Trial (ESWYT) nurseries grown between 1979 and 2021.

**Figure 2.**
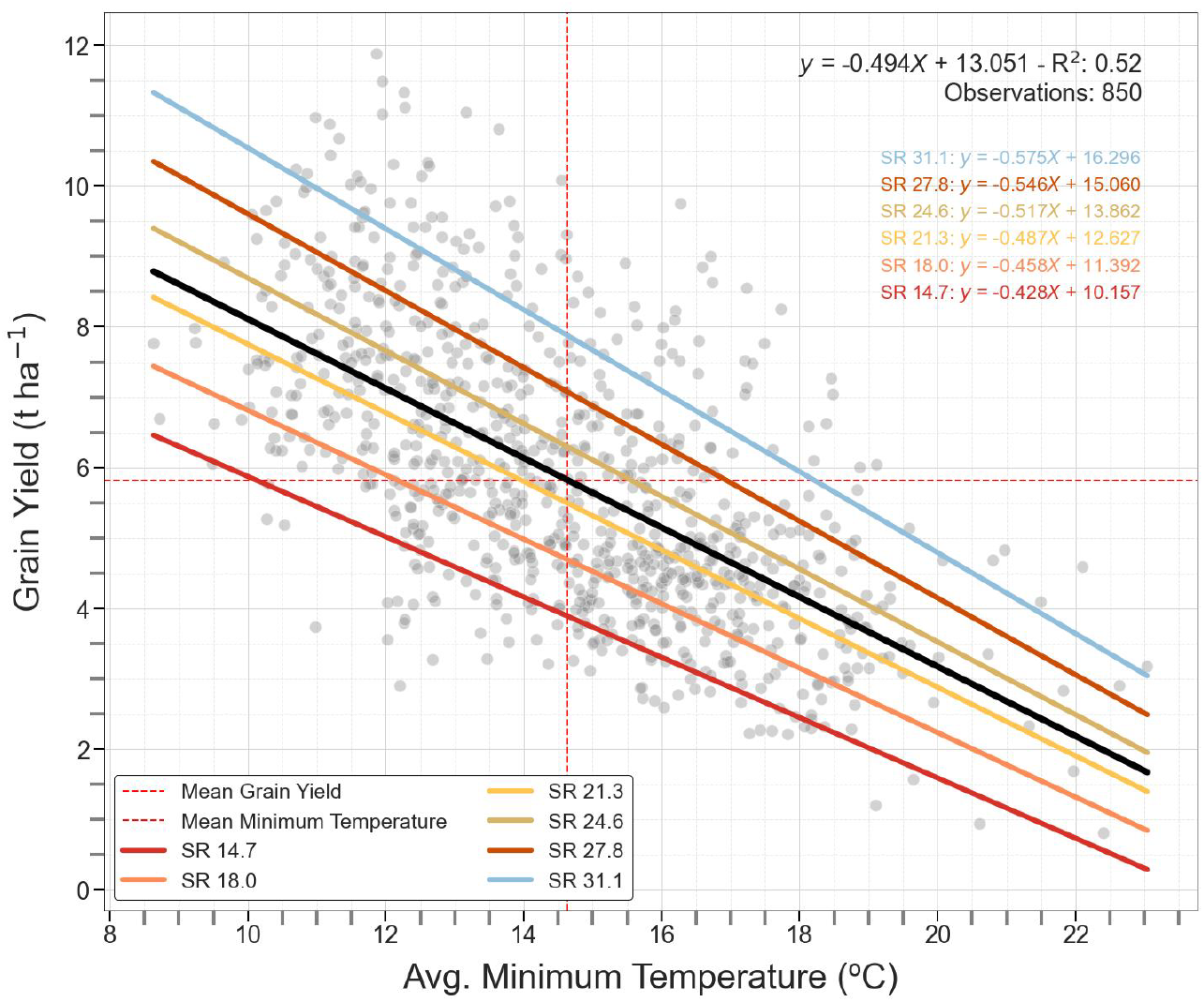
Effect of average daily minimum temperatures and solar radiation during the grain filling period on the average yield obtained at 850 nursery-site-years. The non-significant interaction effects of different levels of solar radiation (SR in MJ/m^2^/day) are overlaid.

**Figure 3.**
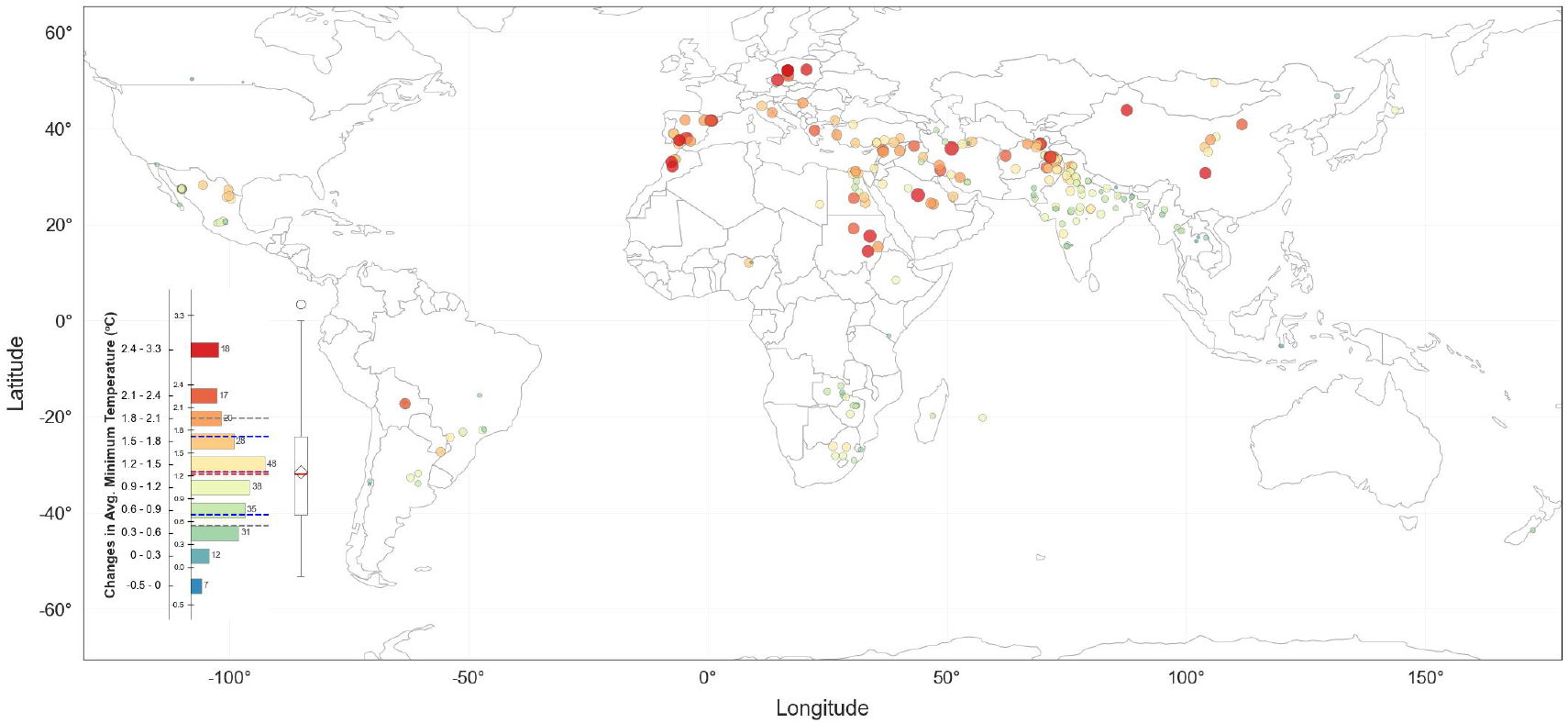
Change in minimum temperature (ºC) at 255 nursery test sites of spring wheat between 1979 and 2021 during the grain filling period.

**Figure 4:**
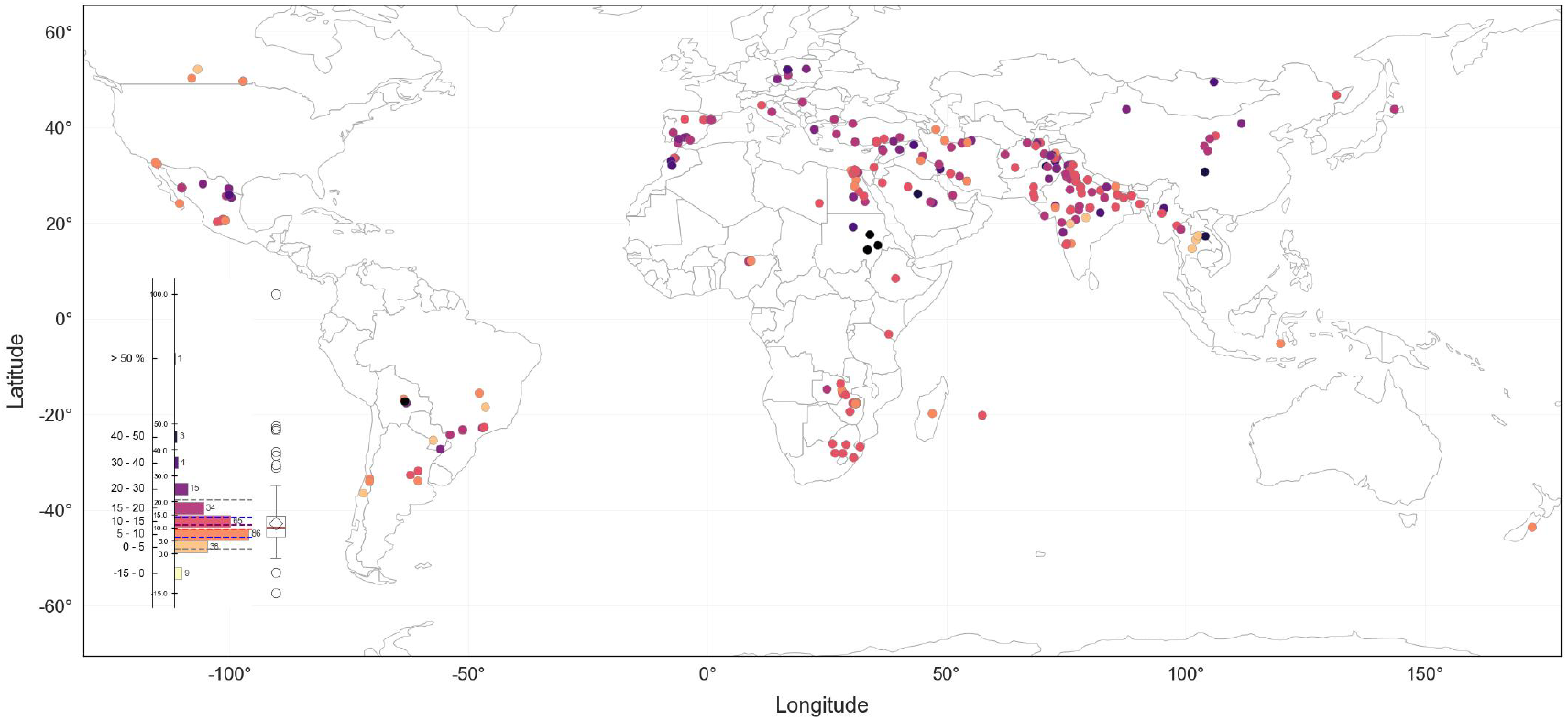
Yield losses (%) of average reported yield caused by changes in daily minimum temperatures during the grain filling period at 255 nursery test sites for spring wheat.

Statistical analyses were conducted with JMP V18.2 (www.jmp.com). First, we identified the variables with the largest prediction power for yield, using the Predictor Screening Platform. For each response, it builds a bootstrap forest model using 100 decision trees. Average daily minimum temperature and solar radiation during the grain filling period had the largest prediction power (Supplementary Fig. 1). We then used the Neural Network platform to model the response and interactions between the 2 parameters. We also used JMP to calculate the variable importance index, which reflects the relative contribution of a factor alone, not in combination with other factors.

In order to quantify and visualize the changes in Tmin during the grain filling period, we calculated the average Tmin for each site-year-occurrence combination, between the start and end day of grain filling for all 42 years. Next, we fit a linear regression of harvest year versus all 42 average Tmin data to estimate the change in Tmin during grain filling at a given location. In a last step, we used the finding of this study that an increase in average Tmin by 1ºC decreases yield by 0.5 t/ha to estimate the yield loss, using the average of the reported yields at a given location as a reference.

## Data availability

The nursery data together with the environmental covariables can be accessed at: https://doi.org/10.71682/10549387

The AgERA5 agrometeorological indicators from 1979 to present derived from reanalysis, can be downloaded from: https://doi.org/10.24381/cds.6c68c9bb

## Acknowledgements

We gratefully acknowledge the support of Digital Transformation and the Climate Action Science Programs of CGIAR, and the International Wheat Yield Partnership (IWYP) funded by the Biotechnology and Biological Research Council of the UK.

## Author contributions

US, MPR and SS conceptualized the study. US conducted the data analysis. US, OA and MPR wrote the initial draft and EG wrote the software for data analysis and visualization. SS was responsible for funding acquisition and project administration. All authors contributed suggestions and reviewed and refined the text.

---

**Supplementary Figure 1:**
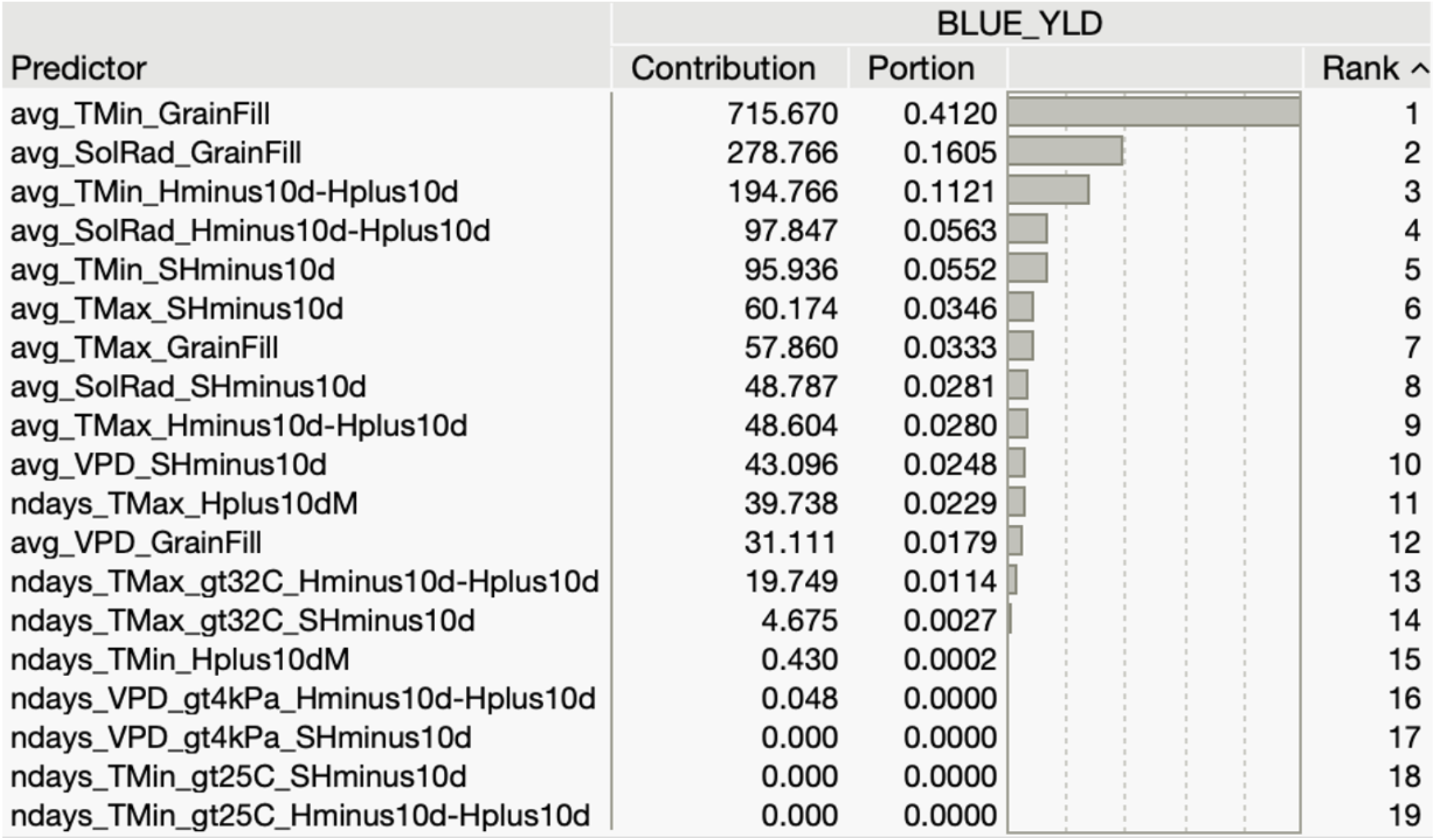
Prediction power of environmental variables to estimate grain yield at 850 site-years. Variables represent three periods: 1) sowing to heading minus 10 days (SHMinus10d), 2) heading minus 10 days to heading plus 10 days (HMinus10d-Hplus10d), and 3) heading plus 10 days to maturity (Hplus10dM). The last period represents grain filling.

**Supplementary Figure 2:**
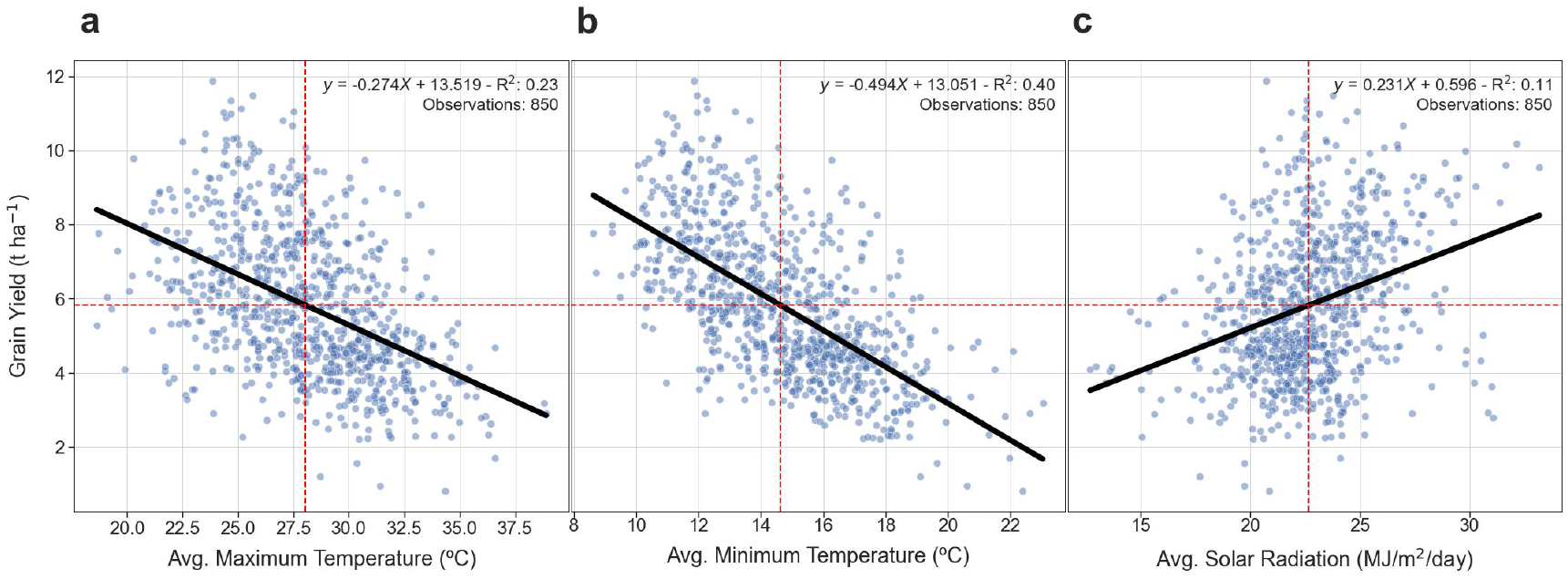
Prediction of wheat yield at 850 site-years as a function of **a)** daily average maximum temperature, **b)** minimum temperature and **c)** solar radiation observed during the grain filling period.

**Supplementary Figure 3:**
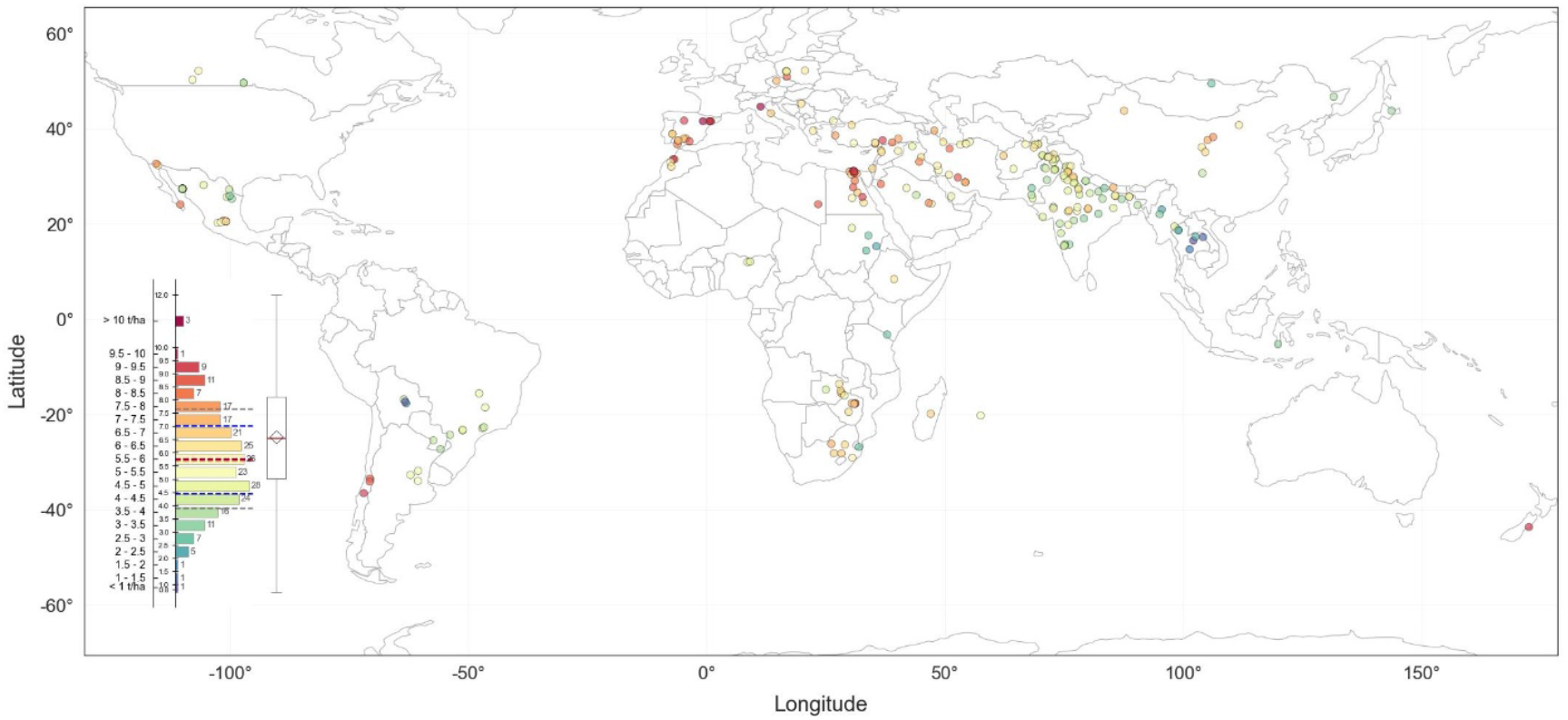
Observed average yield at 255 spring wheat nursery test sites.

**Supplementary Figure 4:**
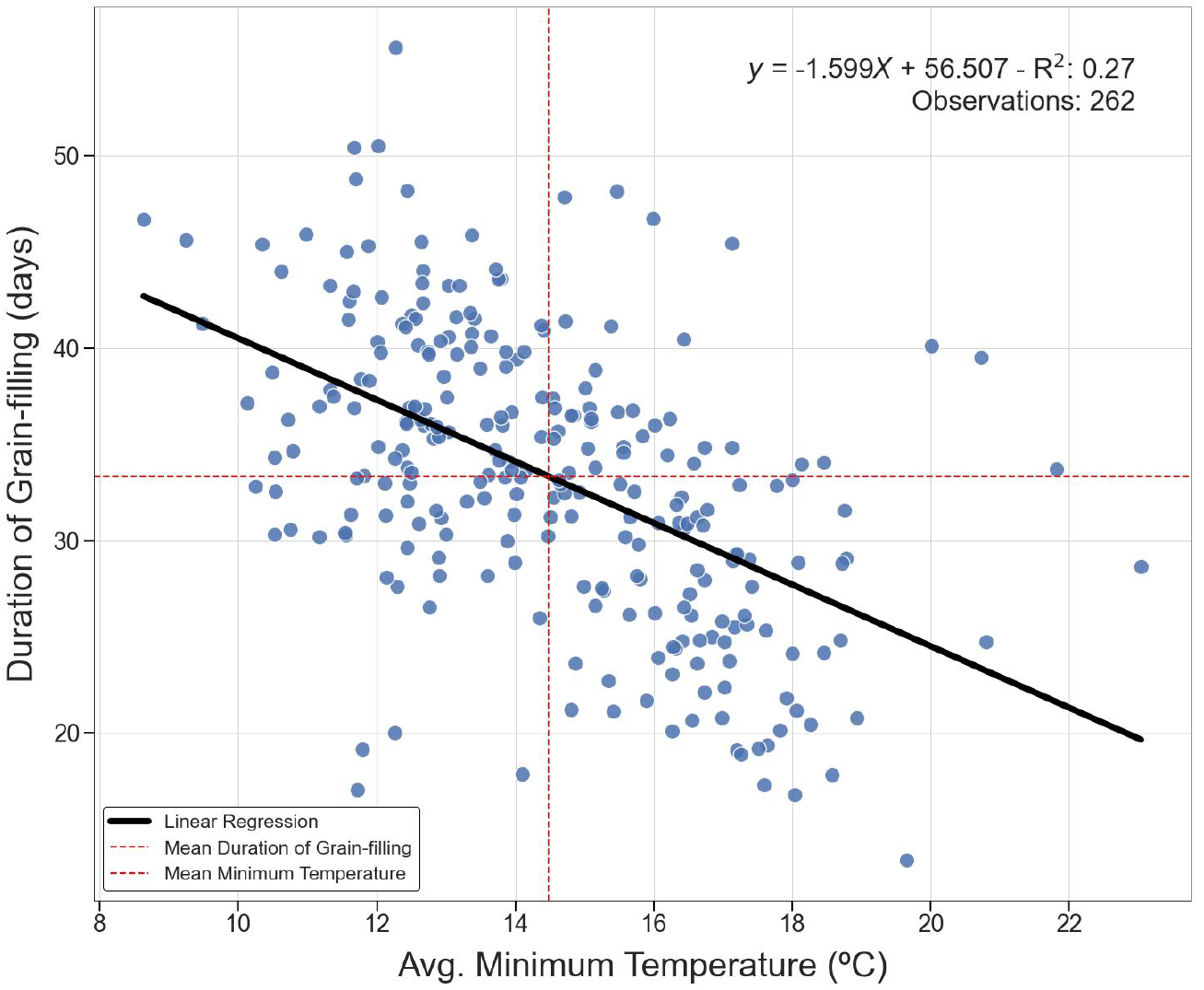
Effect of minimum temperature during grainfilling on the number of reported days between heading and maturity at 262 site-years.

**Supplementary Table 1:**
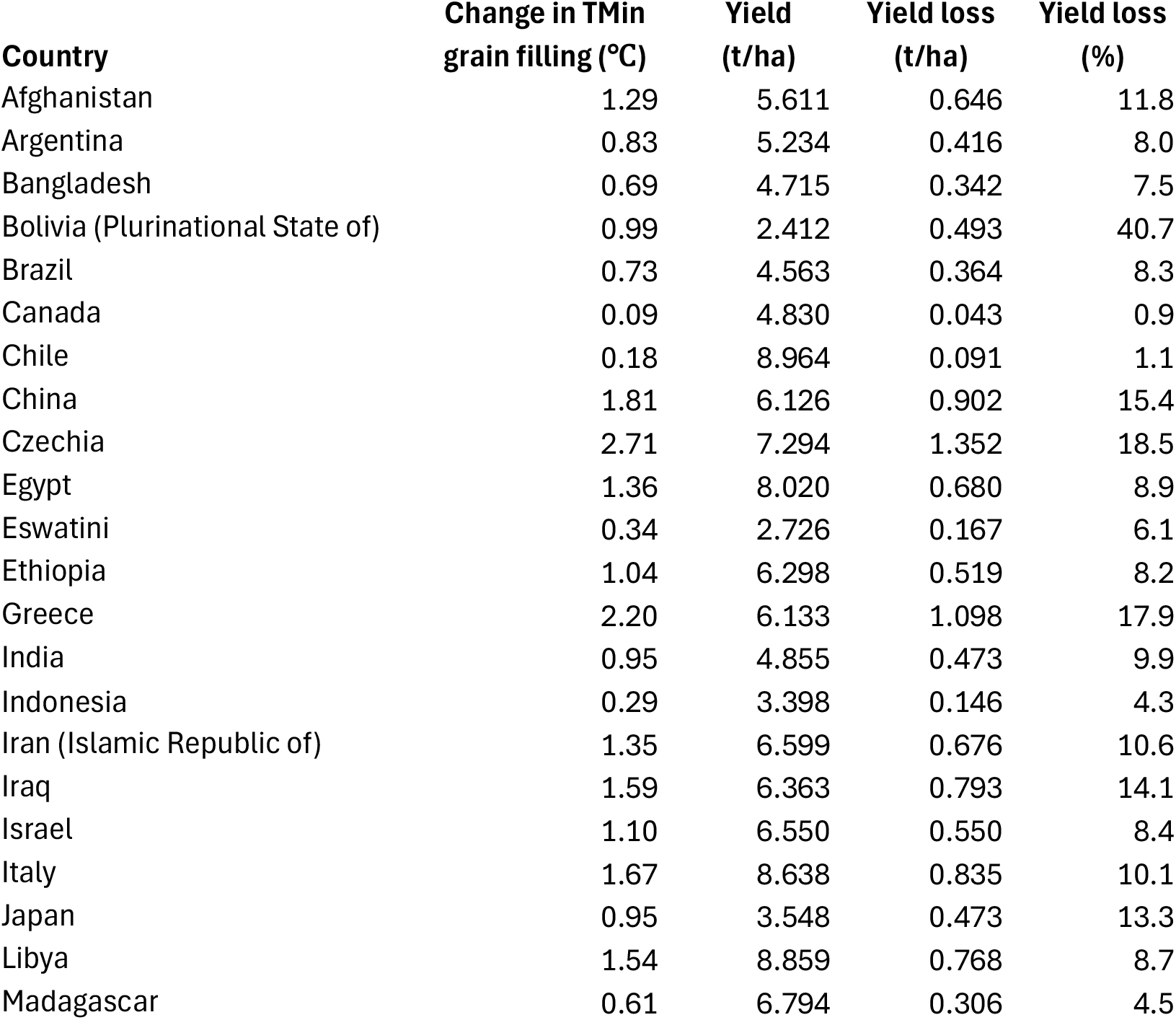
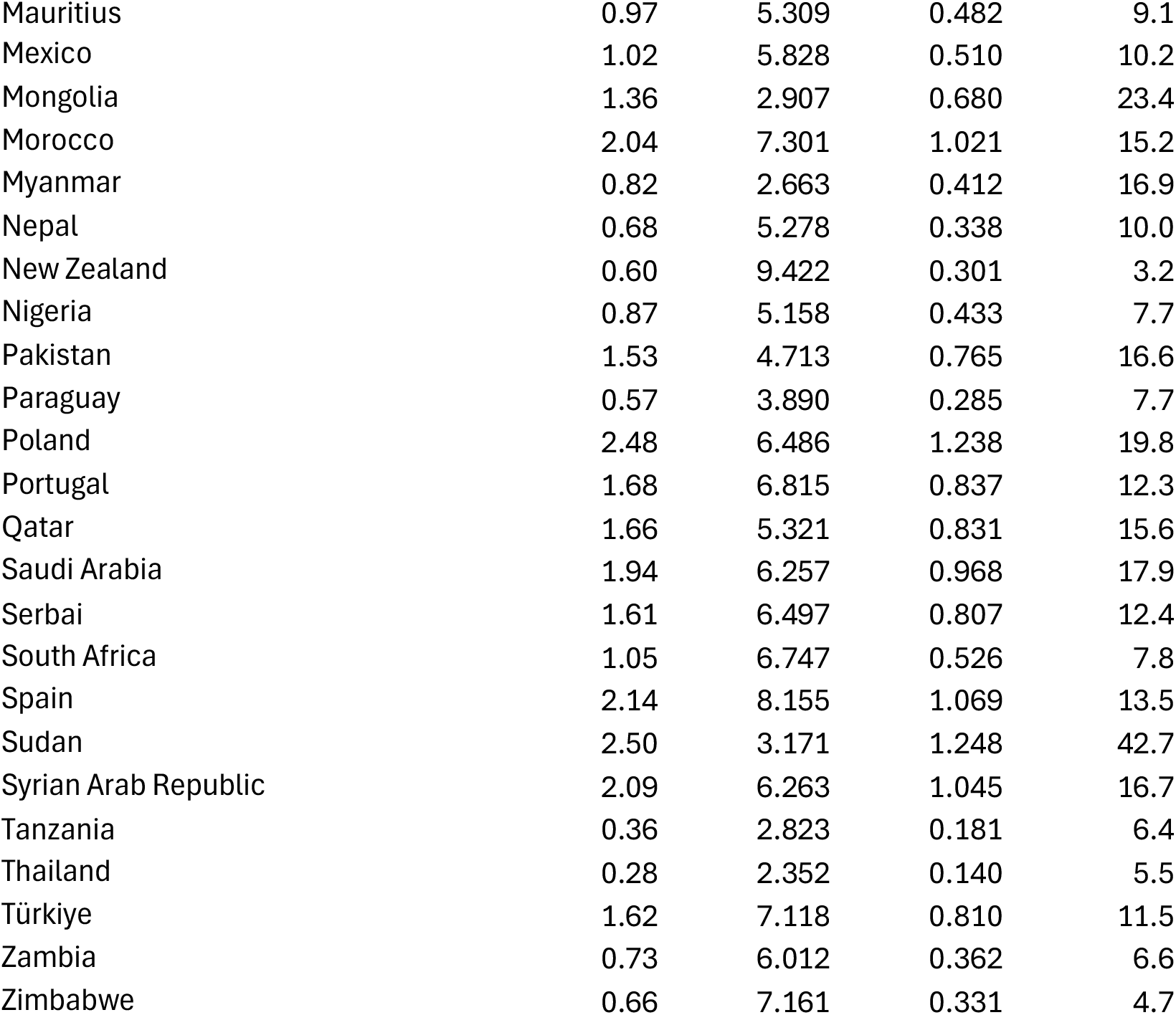
Average change in minimum temperature (Tmin) during grain filling at the ESWYT nurseries test sites. Yield data represent the average nursery yield reported within a country, and yield losses are expressed as a percentage of the reported yield. The data cover 42 years, from 1979 until 2021.

